# Identification of mycobacterial efflux pump inhibiting compounds from *Berberis holstii* Engl. and determination of their mechanism of action

**DOI:** 10.1101/2025.08.15.670474

**Authors:** Ephantus G. Ndirangu, Caroline N. Maina, Edwin K. Murungi, Margaret M. Ng’ang’a, Robi. Chacha, Moses N. Kehongo, Sospeter N. Njeru, Nuhu I. Tukur, Gabriel T. Mashabela, Digby F. Warneh, Paul R. Race, Elizabeth M. Kigondu

## Abstract

*Mycobacterium tuberculosis* (*Mtb*), the causative agent for tuberculosis (TB), is the leading infectious killer of humankind. In 2023, an estimated 10.8 million new cases of TB and 1.25 million deaths were reported globally. Sub-Saharan Africa faces a heavy TB burden, worsened by high HIV-AIDS prevalence and rising drug resistance, making novel anti-TB therapies a pressing priority.

This study investigated the efflux inhibition (EI) activity of compounds characterised from *Berberis holstii* Engl. extracts. Following *in vitro* evaluation of the antimycobacterial activity of aqueous and organic extracts against the non-pathogenic mycobacterial model, *Mycobacterium smegmatis* (*Msm*), and *Mtb*. Thereafter, molecular docking of several compounds identified in chromatographic fractions onto Rv1258c, MmpS5-MmpL5 and Rv2333c *Mtb* efflux pumps (EPs) was undertaken to infer their binding modes and affinities. Subsequently, fractions containing potentially active compounds were tested in combination with spectinomycin (SPEC) and fractional inhibitory concentration index (FICI) values determined. Validation of *Mtb* efflux pump inhibition was performed using CRISPRi knockdown strains of Rv1258c and Rv2333c.

MeOH extracts of the root and the stem bark, and aqueous extracts of the roots, exhibited minimum inhibitory concentration (MIC_99_) values of 906.25μg/ml, 4500 μg/ml and 1875 μg/ml, respectively, against wild-type *Msm*. On the other hand, MeOH extract of the leaves and aqueous extract of the stem bark had MIC_99_ values of 60.43 μg/ml and 1.74 μg/ml respectively, against wild-type *Mtb*. Molecular docking of chillanamine, isoboldine, berberrubine, reticuline, N-methylcoclaurine, thalifoline, apoglaziovine, and orientine onto *Mtb* Rv1258c, MmpS5-MmpL5 and Rv2333c revealed string binding affinities of <-5 kcal/mol. Of the seven fractions containing the bioactive compounds evaluated in checkerboard assays, six were synergistic with SPEC (FICI <0.5), with two fractions lowering the MIC_99_ of SPEC by 8-fold against wild-type *Mtb*.

This study identified phytochemicals with probable *Mtb* efflux pump inhibition, warranting further investigation for potential use in TB combination therapy.

## 1.0 Introduction

Tuberculosis (TB), an infectious disease caused by *Mycobacterium tuberculosis* (*Mtb*), is a leading cause of morbidity and mortality worldwide. In 2023, an estimated 10.8 million new TB cases that resulted and 1.25 million deaths were reported (1). Low- and middle-income countries (LMICs) bear a disproportionate burden of TB which accounts for approximately 95% of total TB-related deaths (2–4). In Sub-Saharan Africa, the deleterious impact of TB is amplified by the vast number of HIV-AIDS cases and the burgeoning antimicrobial resistance (AMR) (5,6). Specifically, the transmission of multidrug-resistant (MDR), extensively drug-resistant (XDR) and totally drug-resistant (TDR) *Mtb* strains necessitates the urgent development of novel anti-TB therapies and potentiators of current agents (7).

Various mechanisms of intrinsic *Mtb* drug resistance including cell wall impenetrability, target modification, drug efflux and over-expression of antimicrobial agents deactivating enzymes have been delineated (8–10). By extruding anti-TB drugs from mycobacterial cells, membrane localised efflux pumps (EPs) diminish the efficacy of these drugs (9,11). To date, numerous *Mtb* EPs, including Rv1819c, Rv1258c, Rv2333c and MmpS5-MmpL5 have been unveiled and characterized to varying degrees (12). Notably, Rv1258c has been linked to the resistance to isoniazid (INH), rifampicin (RIF), spectinomycin (SPEC), fluoroquinolones, tetracycline and clofazimine (Figure **1**) (13), while Rv2333c is associated with resistance to RIF, SPEC and tetracycline (12). MmpS5-MmpL5 is implicated in the resistance to bedaquiline (BDQ), clofazimine and tetracycline (14). Thus, the development of novel *Mtb* efflux inhibitors (EIs) is an attractive strategy for augmenting the efficacy of current anti-TB drugs. Indeed, in TB therapy, synergistic drug combinations are frequently utilised to counter pathogen resistance(15,16).

**Figure 1:**
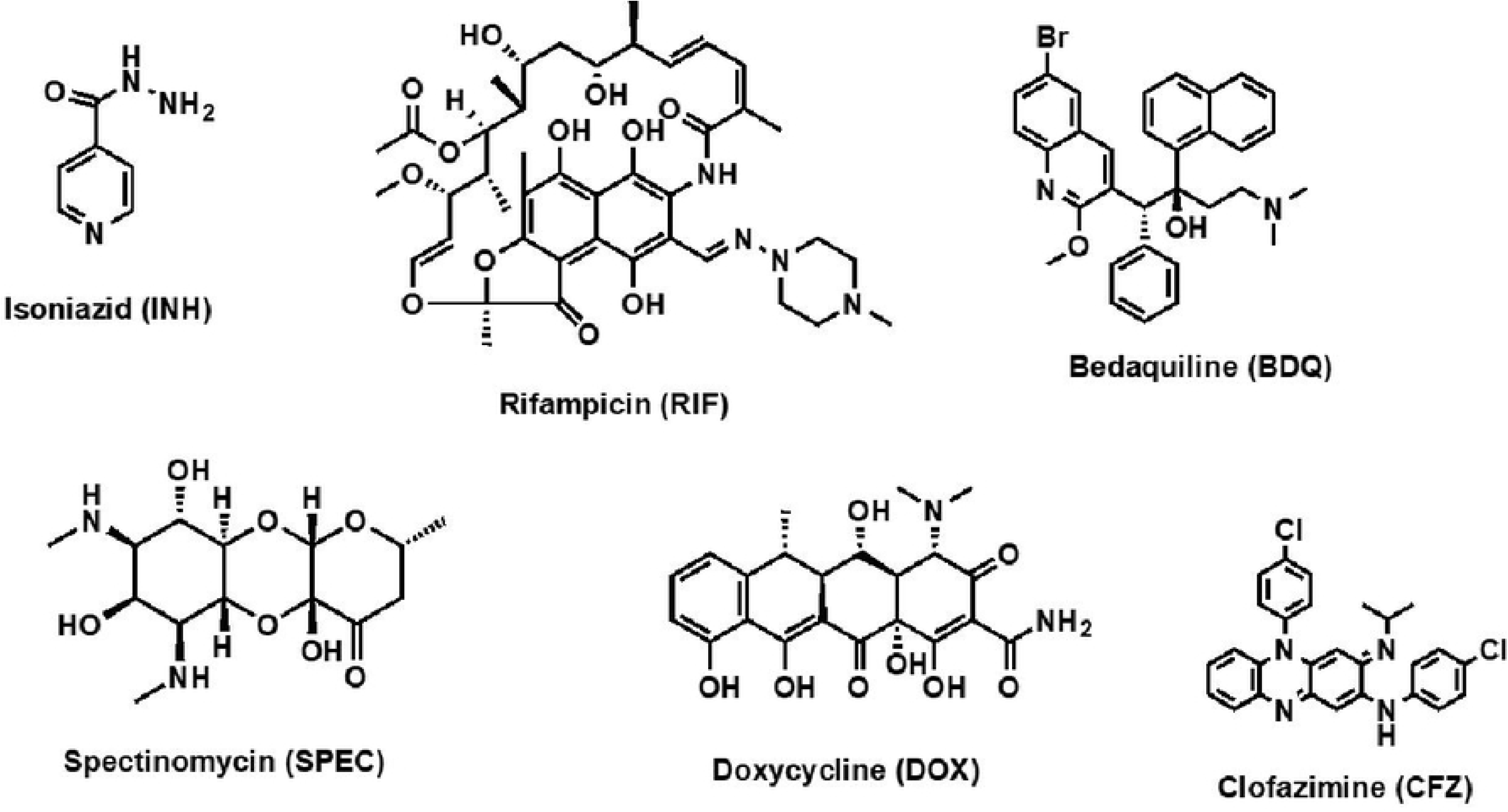
Examples of anti-TB drugs extruded from mycobacterial cells by efflux pumps.

Natural products (phytochemicals) offer a rich source of potential antimicrobials for treating infectious diseases, including those where drug resistance is already established. Importantly, herbal remedies are widely used in LMICs where access to quality healthcare is scant and prohibitively costly for the majority of the populations (17). *Berberis holstii* Engl. (Berberidaceae), commonly known as Holst’s barberry (Figure **2**), is native to seven African countries (Somalia, Kenya, Uganda, Ethiopia, Tanzania, Malawi, and Zambia) and is used for the management of a myriad of diseases including malaria, sexually transmitted diseases, pneumonia, asthma, menorrhagia, and throat infections (18).

**Figure 2:**
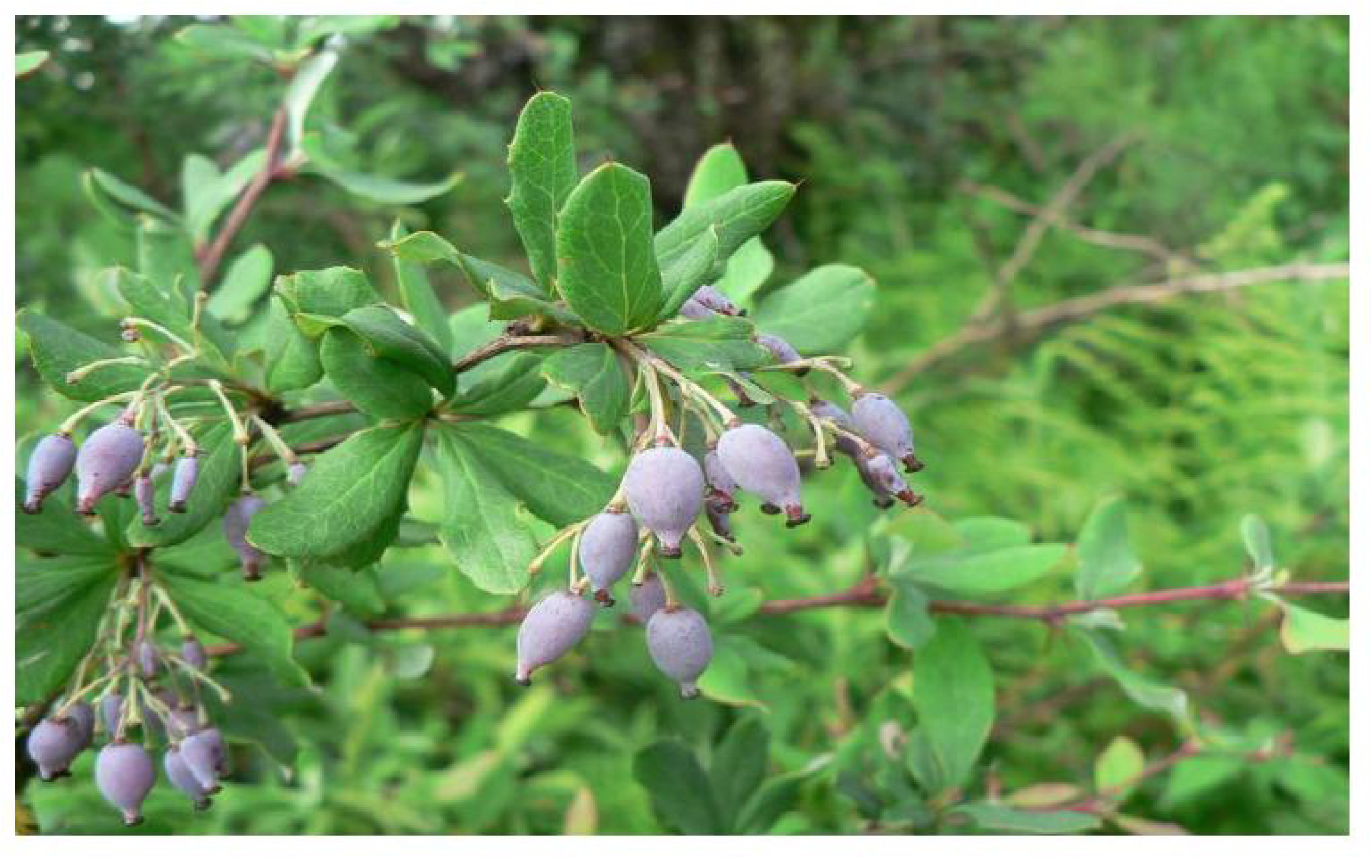
B. holstii plant (http://www.africanplants.senckenberg.de/root/index.php? pageid=78&id=6859)

The plant, a shrub that grows in the open highland forest, the edges and clearings of upland rainforests, and in upland evergreen bushland, grows to a height of approximately 6 m, with 3 cm long spines, short axillary shoots, and oval green berries that turn purple upon ripening, with each fruit containing at most four seeds (18). Isoquinoline alkaloids are the most common bioactive constituents of *Berberis* species whereby berberine is the predominant protoberberine alkaloid (19). Remarkably, it has been reported that berberine and piperine disrupt *Mtb* efEPs) pointing to the probable utility of plant derived natural products as likely *Mtb* EIs (19,20).

In this study, a series of extracts and chromatographic fractions derived from different parts of *B. holstii* were prepared and evaluated for their potential *Mtb* efflux inhibition (EI) activity in combination with RIF and SPEC. RIF, a first line antitubercular agent, exerts its bactericidal activity via high affinity binding to the β-subunit of the DNA-dependent RNA polymerase resulting in transcription disruption (21). The broad-spectrum SPEC inhibits bacteria protein synthesis by blocking translocation of messenger RNA and transfer RNAs on the ribosome (22). RIF and SPEC were selected for this study because of their recognition as substrates for Rv1258c and Rv2333c *Mtb* EPs (12). Although SPEC is not used in the treatment of TB, it was selected for this study due to its relatively narrow spectrum of activity, which is primarily attributed to efflux pump-mediated resistance mechanisms(23). This property makes it a suitable model for demonstrating EI in *Mtb*.

## 2.0 Materials and methods

### 2.1 Collection of plant materials and preparation of extracts

Various *B. holstii* Engl. parts (leaves, stem, and bark) were collected from Cheptongei Forest in Marakwet (coordinates: 0°30’ 50.04’ ’ N, 35° 17’ 21.82” E), identified by a taxonomist, and the associated specimen voucher (CFW/31/1/19/002) deposited at the University of Eldoret, Botany Department Herbarium, in Kenya. Thereafter, the various parts of the plant were air-dried for two weeks and then ground using an electric grinder. The samples were packed in labelled brown paper sleeves and stored in a dry place to prevent compound degradation prior to solvent extraction.

### 2.2 Chemicals

All reagents used during this study were obtained from Sigma-Aldrich and were certified analytical grade. Flat-bottomed Costar 96-well plates were sourced from Merck.

### 2.3 Cells and organisms

For initial extract screening, the non-infectious *Msm* ATCC607 wild-type (WT) was used as a surrogate for *Mtb*. WT *Msm* was obtained as a frozen glycerol stock from the TB laboratory, Centre for Respiratory Diseases Research (CRDR)-KEMRI. Extracts that were found to be active against WT *Msm* were further evaluated against the virulent laboratory strain, *Mtb* H37Rv wild type (WT), at the Molecular Mycobacteriology Research Unit (MMRU), Biology Laboratory, University of Cape Town and at the Center for Virus Centre (CVR), Kenya Medical Research Institute (KEMRI) Nairobi. For validation of EI activity, *Mtb* CRISPRi knockdown strains Rv1258c and Rv2333c, generated by Dr. Gabriel Mashabela at the South African Medical Research Council (SAMRC), Stellenbosch University, were employed. Cytotoxicity assays were performed using African green monkey kidney epithelial cell lines (Vero cells P18), which were seeded from a passaged stock at the Centre of Traditional Medicine and Drug Research (CTMDR), KEMRI.

### 2.4 Solvent extraction

Aqueous extraction was performed by suspending the ground plant material in distilled water at 50 °C and allowing it to stand for 2 hours. The mixture was then filtered, and the resulting filtrate was rapidly frozen using dry ice before being subjected to lyophilization to obtain a solid extract. For organic extracts, ground plant materials were sequentially extracted in the order of solvent polarity, starting from the least to the most polar [n-hexane (HEX), dichloromethane (DCM), ethyl acetate (EtOAc) and methanol (MeOH)]. 200 g of ground plant material was weighed, transferred to a one-litre flask, suspended in 600 mL of n-hexane, and allowed to stand for 48 h. This slurry was then filtered using Whatman filter paper No. 1, with the resulting residue re-extracted using fresh n-hexane for 48 h, followed by filtration. All filtrates were pooled and concentrated in a rotary evaporator (BUCHI Labortechnik AG, 9230 Flawil, Switzerland) under reduced pressure, and at a temperature lower than the solvent’s boiling point. The concentrate was transferred to a pre-weighed labelled vial and the remaining solvent allowed to evaporate at room temperature for 2 weeks. After drying, the remaining residue was sequentially extracted using DCM, EtOAc and MeOH. Dried extracts were subsequently weighed and stored in air-tight vials at 4 °C. For each extract, the percentage yield was calculated.

### 2.5 *In vitro* antimycobacterial screening

Extracts were assayed for antimycobacterial activity, with minimum inhibitory concentration (MIC_99_) values determined using the Microplate Alamar Blue Assay (MABA) format (24–27).

#### 2.5.1 Preparation of plant extracts and reference stock solutions

Stock solutions comprised 50 mg of dried plant extract dissolved in 1000 μL of dimethyl sulfoxide (DMSO), mixed by vortexing for 10 min. A stock concentration (0.2 mg/mL) of the reference drug (RIF) was prepared by dissolving 1 mg of compound in 1 mL of DMSO. Total plant extracts and RIF stock solutions were sterilised by filtration using a polyethersulfone (PES) 0.22 μm syringe followed by gamma irradiation and stored at 4 °C for antimycobacterial activity assays.

#### 2.5.2 Preparation of Middlebrook 7H9 broth media

Middlebrook 7H9 media was prepared as follows: 4.7 g of Middlebrook 7H9 powder was dissolved in 50 mL of distilled water and the volume adjusted to 900 mL in a Schott bottle. 4 mL of glycerol was subsequently added to the mixture, which was then autoclaved at 121 °C for 10-15 min, followed by cooling to 50-55 °C. To this solution, 100 mL of Middlebrook Oleic Acid Albumin Dextrose Catalase (OADC) enrichment was aseptically added, followed by the addition of 4 mL of 20% Tween 80 (prepared as 20 mL of Tween 80 in 80 mL of distilled water) to prevent clumping. The media was stored at 2-8 °C, with a 50 mL sample incubated at 37 °C for 7 days to ensure sterility prior to use.

#### 2.5.3 Preparation of Mycobacterium smegmatis (*Msm*) and *M. tuberculosis* (*Mtb*) culture

Frozen glycerol stocks were pre-cultured for 24 h at 37 °C without shaking. Upon attainment of an optical density (OD_600nm_) of 0.5-0.7, cultures were diluted to an OD_600nm_ of 0.001 in filter-sterilized 7H9 broth media.

#### 2.5.4 Microplate Alamar Blue (MABA) Assays

MABA assays were performed as described previously (24–26,28) with minor modifications. Two-fold dilution in a 96-well plate was used to determine the MIC_99_. 50 µL of media was added to each well of the plate, with an additional 50 µL added to the wells in A2 to H2, to which 5 µL of plant extract was dispensed in triplicate at an initial concentration of 50 mg/mL. The cells from A1-to-H1 and A12-to-H12 wells were designated as minimum inhibition and maximum inhibition controls, respectively. For the two-fold serial dilution, 50 µL of the liquid (plant extract + broth media) was taken from A2 to H2 wells and serially diluted to A11-to-A11 wells and 50 µL discarded. A 50 µL aliquot of pre-grown *Msm* ATCC607 wild type (WT) *Msm* or H37Rv wild type (WT) *Mtb* inoculum was added to all wells. Microplates were carefully sealed and incubated for 48 h for *Msm* and 13 days for *Mtb* at 37 °C. Thereafter, 20 µL of 0.01% freshly prepared resazurin dye in sterile distilled water was added to each well of the plate, which was subsequently incubated for a further 8 h for *Msm* and 24 h for *Mtb*. A colour change from blue to pink indicated bacterial growth. MIC_99_ was defined as the lowest concentration of drug that inhibits the visible growth of the mycobacteria (blue color). RIF was used as a reference standard drug in the assays involving extracts.

#### 2.5.5 Checkerboard assay

A standard two-dimensional (2-D) checkerboard assay of each extract/fraction with either SPEC or RIF performed in a 96-well plate was used to ascertain whether the compound interactions between the “drug” partners was additive, synergistic or antagonistic. The fractional inhibitory concentration index (FICI) for each compound was calculated as detailed previously in (29) as follows: FIC_A_ = (MIC of compound A in the presence of compound B)/(MIC of compound A alone), where FIC_A_ is the fractional inhibitory concentration of compound A. The FIC for compound B was calculated in the same manner. FICI was calculated as the sum of FIC_A_ and FIC_B_. Synergy was defined as the observation of an FICI value of ≤0.5, antagonism by an FICI value of >4.0, and no interaction by an FICI value of 0.5 - 4.0 (30).

The efflux EI activity of the natural product-derived EPIs was validated using CRISPRi knockdown *Mtb* strains Rv1258c and Rv2333c, generously donated by Dr. Gabriel Mashabela based at the South African Medical Research Council (SAMRC), Stellenbosch University. The mutant freezer stocks were revived in 7H9 medium supplemented with 25 μg/ml kanamycin and incubated at 37 °C in vented T25 cell culture flasks without shaking, until an OD_600nm_ of 0.5–0.7 was reached. Upon reaching the desired OD, the cultures were sub-cultured into fresh medium and diluted to the same OD_600nm_. Anhydrotetracycline (ATc) was then added to a final concentration of 200 ng/ml as an inducer for regulating gene expression in knockdown *Mtb* strains by binding to the TetR repressor protein, thereby alleviating TetR-mediated transcriptional repression of target genes (31).

### 2.6 Cytotoxicity assay

To assess the cytotoxicity of plant extracts against mammalian cells, 3-(4,5-dimethylthiazol-2-yl)-2,5-diphenyltetrazolium bromide (MTT )assays were conducted, monitoring the viability of Vero cells in the presence of plant extracts (32). Plant extracts were prepared as 20 mg/mL stock solutions in 10% DMSO. Stock solutions were serially diluted 10-fold in a complete medium, to generate six concentrations. Following incubation with test compounds for 48 h, cells were treated with MTT and subsequently incubated at 37 °C for an additional 4 h. RIF was used as a positive control, with media. All assays were performed in triplicate. IC_50_ values were derived from dose-response curves using non-linear dose-response curve fitting analysis performed using GraphPad Prism v.4 software (33).

### 2.7 Fractionation of plant extracts

Fractionation of non-cytotoxic extracts that showed activity against WT *Msm* or WT *Mtb* was performed as follows. Plant extracts were ground to a fine powder, mixed with silica gel of equal mass, extraction solvent added, and the mixture stirred to dissolve the extract. Following the mixing, the solvent was evaporated using a rotary evaporator to dryness. The solid mixture was again ground to a powder. A column, loaded with a mixture of hexane and silica gel (pore size 60 Å, 230-400 mesh particle size, 40-63 μm) was prepared and allowed to stand for 2 h. The powdered mixture of the plant extract and the silica gel was carefully loaded onto the column and levelled (0.04-0.063 mm, 120 g). Column chromatography using HEX, DCM, EtOAc and MeOH as the mobile phase was performed. Column elution was commenced using 100% hexane. Fractions of 100-150 mL each were collected based on spots observed on thin layer chromatography (TLC). The polarity of the solvent system was gradually increased using DCM, EtOAc and MeOH, sequentially. Fractions were concentrated under reduced pressure using a rotary evaporator and subsequently pooled based on similarities in their TLC profiles. Spots visualization was performed under ultraviolet lamp, Spectroline Model EN-280C/FE (UV, 254 and 366 nm) followed by staining with vanillin–sulfuric acid reagent (prepared by dissolving 15 g of vanillin in 250 mL of methanol and adding 2.5 mL of concentrated sulfuric acid), and then gently heating the plates to enhance spot development. Preparative thin layer chromatography (PTLC) was used to separate out semi-pure compounds with more than one spot. Semi-dried samples were allowed to stand at room temperature until all the solvent had evaporated. The samples were then placed in vials, sealed and stored at 4 °C.

### 2.8 Liquid chromatography coupled mass spectrometry (LC-MS) analysis

LC-MS analysis was performed using a Waters G2-XS quadruple time-of-flight (QTof), incorporating a Waters Acquity UPLC system equipped with an autosampler. Chromatographic separation was performed on a Supelco Purospher STAR RP-18 end-capped UHPLC column (100 mm × 2 mm, particle size 2 µm) at 30-40 °C. The mobile phase consisted of water (A) and acetonitrile (B) with a flow rate of 0.25 mL/min. The gradient elution profile used was: a linear gradient increasing from 5% to 95% B (6 min) and 95% B (4 min). The mass spectrometer was operated in negative electrospray ionisation (ESI) mode, with the heating source temperature set at 250 °C, cone voltage at 40 V, desolvation temperature at 350 °C, and desolvation and cone gas flow rate at 700 L/min and 50 L/min, respectively. A mass range from 40-1200 m/z was scanned at a scan rate of 5 ns. One milligram of each sample was dissolved in 1 mL of the solvent used to extract the sample and 10 µL injected into the LC-MS system for analysis. Data from the mass spectra was converted into mzML files using MSConvert software. mzML files were interpreted using MZmine software which was able to display the chromatogram and fragmentation patterns of the compounds (34). The fragmentation pattern and the molar masses of the compounds given by the MZmine software were compared with those reported in literature and the NLM (National Library of Medicine), lotus, and GNPS (Global Natural Product Social Molecular Networking) databases to annotate the compounds identified in the extracts (35).

### 2.9 *In silico* docking

Molecular docking was performed using the Maestro graphical interface of the Schrodinger Suite software package (Schrödinger, LLC, NY, 2019-2) (36). The suite provides various tools for ligand preparation, protein preparation, active site identification and docking (37).

#### 2.9.1 Ligand preparation

The 2-D compounds in SDF format obtained from PubChem were prepared for docking using LigPrep. The Epik module was used for generating the best possible ionization states and tautomer at a pH of 7 (±) and OPLS4 force field chosen (38). A maximum of 32 stereoisomers were generated for each compound.

#### 2.9.2 Protein preparation

Homology models of the *Mtb* proteins of interest (Rv1258c, MmpS5-MmpL5, and Rv2333c) were prepared for docking using the Protein Preparation wizard. The CCD database was used to allocate bonding orders (39). Zero-order bond metals and disulfide bonds were created. Missing side chains and loops were inferred using the Prime module while hetero states were generated using Epik. PROPKA was used to optimise hydrogen bond assignments. Waters 5.0 Å of heteroatoms were removed and the structure was minimized using OPLS4 force field.

#### 2.9.3 Active site prediction and GLIDE grid generation

The SiteMap tool was used to predict the models’ active sites with a maximum of 5 sites generated. Active sites with a druggability score and site score of greater than 7.0 were chosen for docking (39). The Receptor Grid Generation wizard was used to generate the grid files for the selected active sites.

#### 2.9.4 Molecular docking

The Ligand Docking tool was used for docking ligands onto potential targets. The default scaling factor of 0.80 and partial charge cutoff of 0.15 were set as the parameters for the van der Waals radii scaling. Docking was undertaken in the extra precision (XP) mode and Epik state penalties were considered in the final docking score. Thereafter, the docking score function was employed in the ranking of the docking poses of each ligand. A docking score of -5.0 kcal/mol or lower was assumed to infer robust ligand binding capable of disrupting the target function (40).

### 2.10 Data management and analysis

Identification of compounds was performed by comparing their chromatograms and spectra with those of known compounds contained in the NLM, Lotus, and GNPS databases. ChemDraw software (Ultra 12.0) was used to render the structures of the compounds identified. MZmine and MSconvert were used for the deconvolution of chromatograms and interpretation of the mass fragmentation patterns. In order to perform appropriate multiple comparison tests, analysis of variance (ANOVA) was used to test the significance of differences between mean results with p < 0.05 values deemed as statistically significant.

## 3.0 Results

### 3.1 Extraction of *B. holstii* plant parts with solvents

Overall, this study aimed to determine whether *B. holstii* secondary metabolites can potentiate anti-TB drugs, particularly those targeting reported resistant *Mtb* strains, through disruption of EP activity. From the aqueous and organic extracts generated from individual plant parts, the yields resulting from the extracts were determined based on the mass of recovered material. Aqueous extracts had the highest yields, followed by MeOH extracts, while DCM, and HEX gave relatively low yields (Table **1**).

**Table 1.** Extraction yields of *B. holstii* extracts.

| Plant part | % Yield |  |  |  |  |
| --- | --- | --- | --- | --- | --- |
|  | HEX | DCM | EtOAc | MeOH | H<sub>2</sub>O |
| Leaves | 0.3 | 2.5 | 2.4 | 4.5 | 6.1 |
| Stem bark | 0.2 | 0.3 | 0.4 | 4.6 | 6.3 |
| Root bark | 0.4 | 0.2 | 0.2 | 5.3 | 7.2 |

HEX - hexane, DCM - dichloromethane, EtOAc - ethyl acetate, MeOH - methanol, H<sub>2</sub>O - water

### 3.2 Activity of *B. holstii* extracts against WT *Msm* and WT *Mtb*

All the extracts (Table **1**) were tested for antimicrobial activity against WT *Msm* and WT *Mtb*. The overall results show that the tested extracts were more active against WT *Mtb* than the WT *Msm* based on their MIC_99_ (Table **2**). Specifically, all leaf extracts had MIC_99_ of >5000.00 μg/ml in *Msm*, and MIC_99_ of >125.00 μg/ml in WT *Mtb* (except the methanolic extract, which gave an MIC_99_ of 60.43 μg/ml). All stem bark extracts had MIC_99_ of >5000.00 μg/ml against WT *Msm* (except methanolic extract with MIC_99_ of 4500.00 μg/ml), and MIC_99_ of >125.00 μg/ml in *Mtb* (except the water extract with an MIC_99_ of 1.73 μg/ml). Furthermore, the root bark hexane and DCM extracts had MIC_99_ of >5000.00 μg/ml against WT *Msm*, while the methanolic extract had an MIC_99_ of 906.25, and the water extract had an MIC_99_ of 1875 μg/ml against the same strain. On the other hand, all extracts had an MIC_99_ of >125.00 μg/ml in WT *Mtb*. As expected, the positive control, RIF gave the highest activity with a MIC_99_ of 15.10 μg/ml in WT *Msm*, and MIC_99_ of 0.01 μg/ml in WT *Mtb* (Table **2**). Although the extracts exhibited lower MIC₉₉ values than the control, as expected, they likely contain multiple compounds, some of which may possess strong activity but are present in relatively low concentrations(41).

**Table 2.**
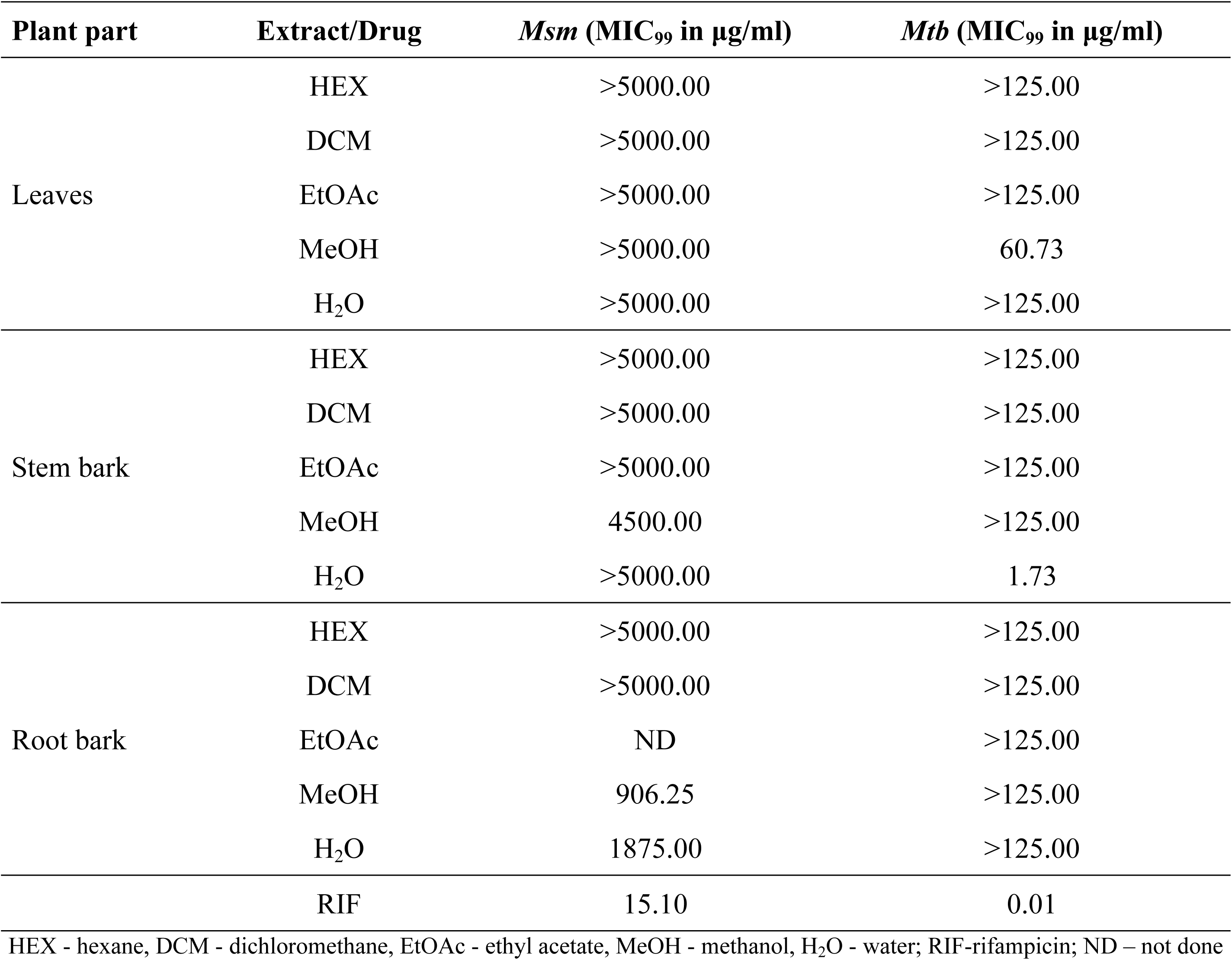
Antimycobacterial activity of *B. holstii* extracts.

| Plant part | Extract/Drug | *Msm* (MIC<sub>99</sub> in µg/ml) | *Mtb* (MIC<sub>99</sub> in µg/ml) |
| --- | --- | --- | --- |
| Leaves | HEX | >5000.00 | >125.00 |
|  | DCM | >5000.00 | >125.00 |
|  | EtOAc | >5000.00 | >125.00 |
|  | MeOH | >5000.00 | 60.73 |
|  | H<sub>2</sub>O | >5000.00 | >125.00 |
| Stem bark | HEX | >5000.00 | >125.00 |
|  | DCM | >5000.00 | >125.00 |
|  | EtOAc | >5000.00 | >125.00 |
|  | MeOH | 4500.00 | >125.00 |
|  | H<sub>2</sub>O | >5000.00 | 1.73 |
| Root bark | HEX | >5000.00 | >125.00 |
|  | DCM | >5000.00 | >125.00 |
|  | EtOAc | ND | >125.00 |
|  | MeOH | 906.25 | >125.00 |
|  | H<sub>2</sub>O | 1875.00 | >125.00 |
|  | RIF | 15.10 | 0.01 |

HEX - hexane, DCM - dichloromethane, EtOAc - ethyl acetate, MeOH - methanol, H<sub>2</sub>O - water; RIF-rifampicin; ND – not done

### 3.3 Cytotoxicity the active extracts of *B. holstii* extracts

After observing antimycobacterial activity in some extracts, notably methanolic extract of the leaves as well as methanolic and water extracts of the stem and the root barks, the safety levels of these select extracts were evaluated using *in vitro* Vero cell line model. Interestingly, all tested extracts were within the acceptable toxicity thresholds (CC_50_ of 201 to 500 μg/ml weakly toxic) (42–45) and thus considered to be safe (Table **3**).

**Table 3.** Cytotoxicity of the active extracts of *B. holstii* extracts.

| Plant part | Extract/Drug | Cytotoxicity ($CC_{50}$) ($\mu\text{g/ml}$) |
| --- | --- | --- |
| Leaf | MeOH | $227.1 \pm 42.8$ |
| Stem bark | MeOH | $432.8 \pm 47.6$ |
| | H<sub>2</sub>O | $359.8 \pm 85.4$ |
| Root bark | MeOH | $226.8 \pm 50.2$ |
| | H<sub>2</sub>O | $487.2 \pm 40.9$ |
| Positive Control | RIF | $246.5 \pm 34.0$ |

RIF-rifampicin; MeOH - methanol, H<sub>2</sub>O – water; $CC_{50} < 20 \mu\text{g/mL}$ highly cytotoxic, $CC_{50}$ 21 to 200 $\mu\text{g/mL}$ moderately cytotoxic, $CC_{50}$ 201 to 500 $\mu\text{g/mL}$ weakly cytotoxic, and $CC_{50} > 500 \mu\text{g/mL}$ noncytotoxic

### 3.4 Checkerboard assays of the active extracts of *B. holstii* against *Msm*

Given the promising antimycobacterial activity and safety of the tested extracts, which is an indication of selective bioactivity, it was important to investigate whether the extracts could reduce the MIC of known antimycobacterial drugs. Consequently, checkerboard assays were conducted. Interestingly, aqueous and MeOH extracts of the root bark displayed synergism (FICI value of ≤0.5) with RIF, while MeOH extract of the stem bark showed an additive effect (FICI value of 0.5 to 4) (28) (Table **4**).

**Table 4.** Checkerboard assay results for the total extract of *B. holstii* in combination with RIF against WT *Msm*.

| RIF/extract | MIC<sub>99</sub> (µg/ml)<br>singly | MIC<sub>99</sub> (µg/ml)<br>combination | MIC<sub>99</sub> (µg/ml)<br>fold reduction<br>of RIF | FIC | FICI |
| --- | --- | --- | --- | --- | --- |
| RIF | 13.0±4.5 | 0.8±0.3 | 16 | 0.0625 | 0.125 |
| Rb-MeOH extract | 755.2±261.6 | 47.2±16.4 |  | 0.0625 |  |
| RIF | 20.83±9.02 | 1.3±0.6 | 16 | 0.0625 | 0.1875 |
| Rb-H<sub>2</sub>O extract | 2500.0±1082.5 | 312.5±132.3 |  | 0.125 |  |
| RIF | 10.4±4.5 | 2.6±1.1 | 4 | 0.25 | 0.7500 |
| Sb-MeOH extract | 3000.0±1299.0 | 1500.0±649.5 |  | 0.5 |  |
| RIF | 18.2±11.9 | 9.1±6.0 | 2 | 0.5 | N/A |
| Sb-DCM extract | >5000 | >5000 |  | N/A |  |

Rb – root bark, Sb – stem bark; RIF - rifampicin; MeOH - methanol, H<sub>2</sub>O - water; N/A- not applicable; FIC- fraction inhibitory concentration index; FICI-fraction inhibitory concentration index

### 3.5 Fractionation and characterization

Many *Berberis* species have been reported to contain berberine, a compound known to inhibit EPs in *Mtb* (19,20). However, there is currently no literature on the compounds isolated from *B. holstii*. Therefore, it was important to identify the compounds present in *B. holstii*, as it may contain other potential EIs besides berberine. Total MeOH extracts of the root bark, stem bark, and leaves of *B. holstii* were selected for fractionation based on their ability to lower the MIC_99_ of RIF by ≥ 4-fold. Extracts were subjected to column chromatography and separated into 8, 9, and 4 fractions of root bark, stem bark, and leaves, respectively. The fractions were subsequently characterized using LC-MS (Table **5**, Supplementary Table **S1,** and Supplementary Table **S2**). For compound identification, the molar masses and fragmentation patterns of individual compounds were compared to those reported in literature and those contained in the NLM, Lotus, and GNPS repositories. The compounds shown in (Table **5**, Supplementary Table **S1** and Supplementary Table **S2)** have been reported in other *Berberis* species (46).

**Table 5.**
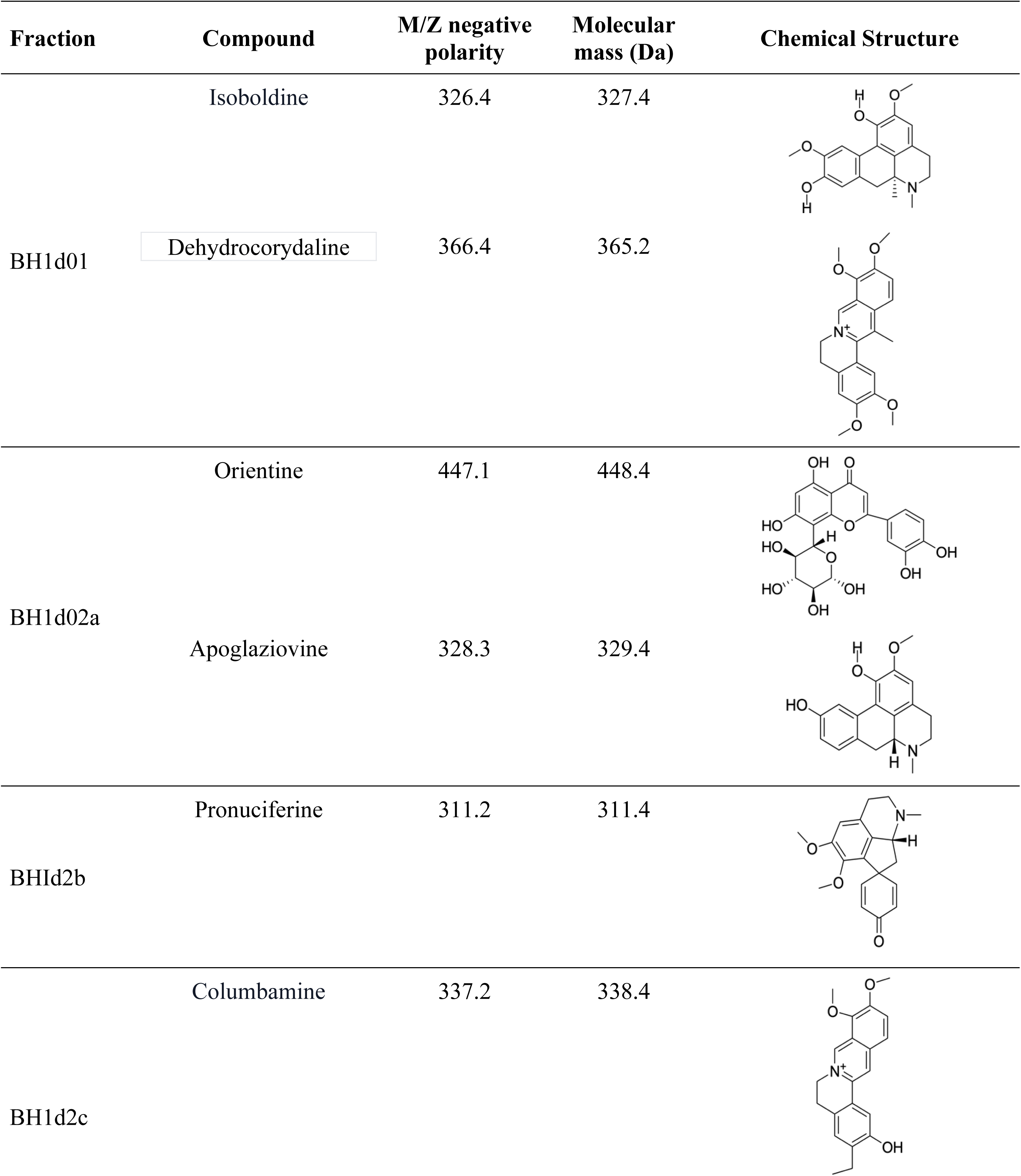
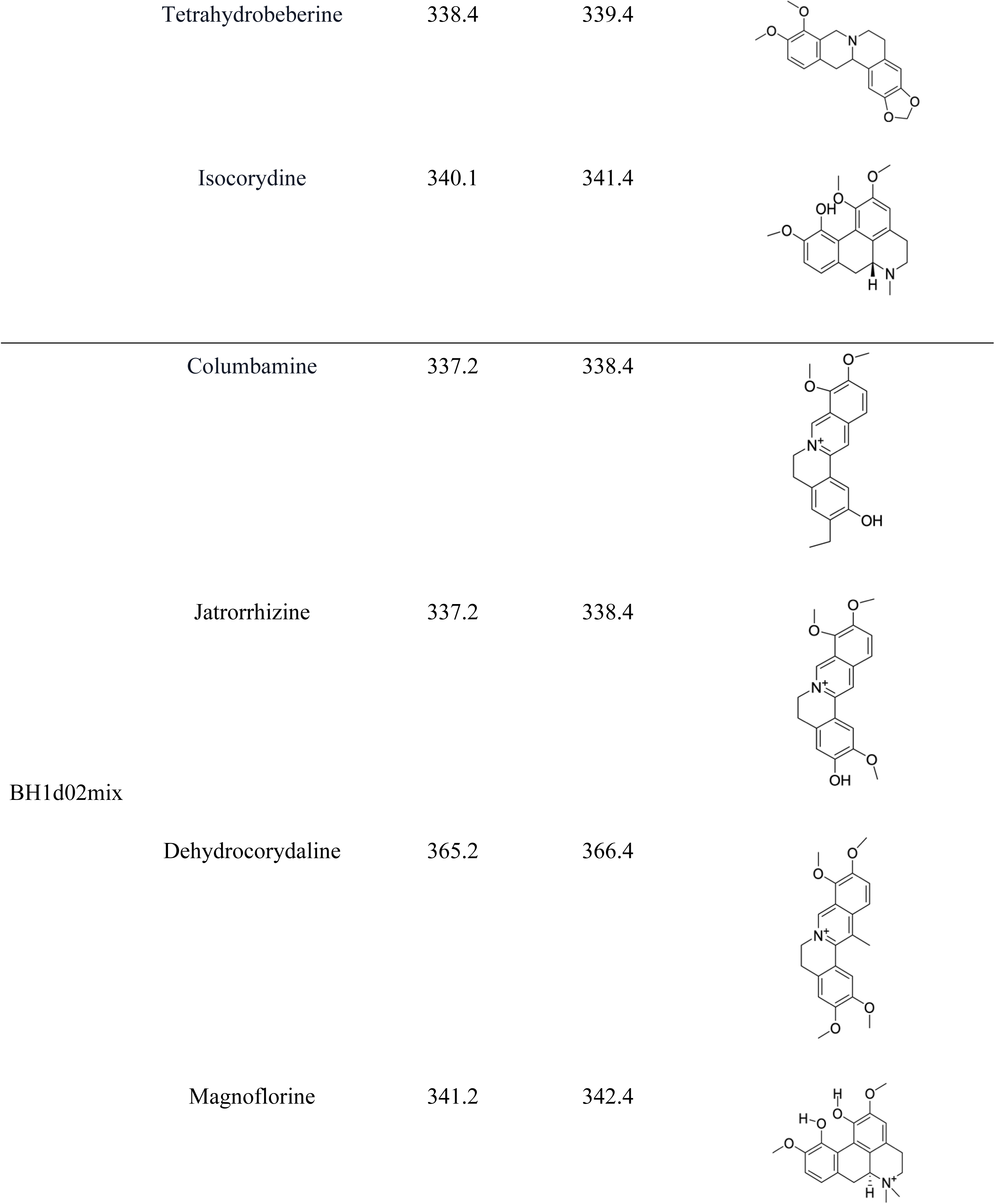
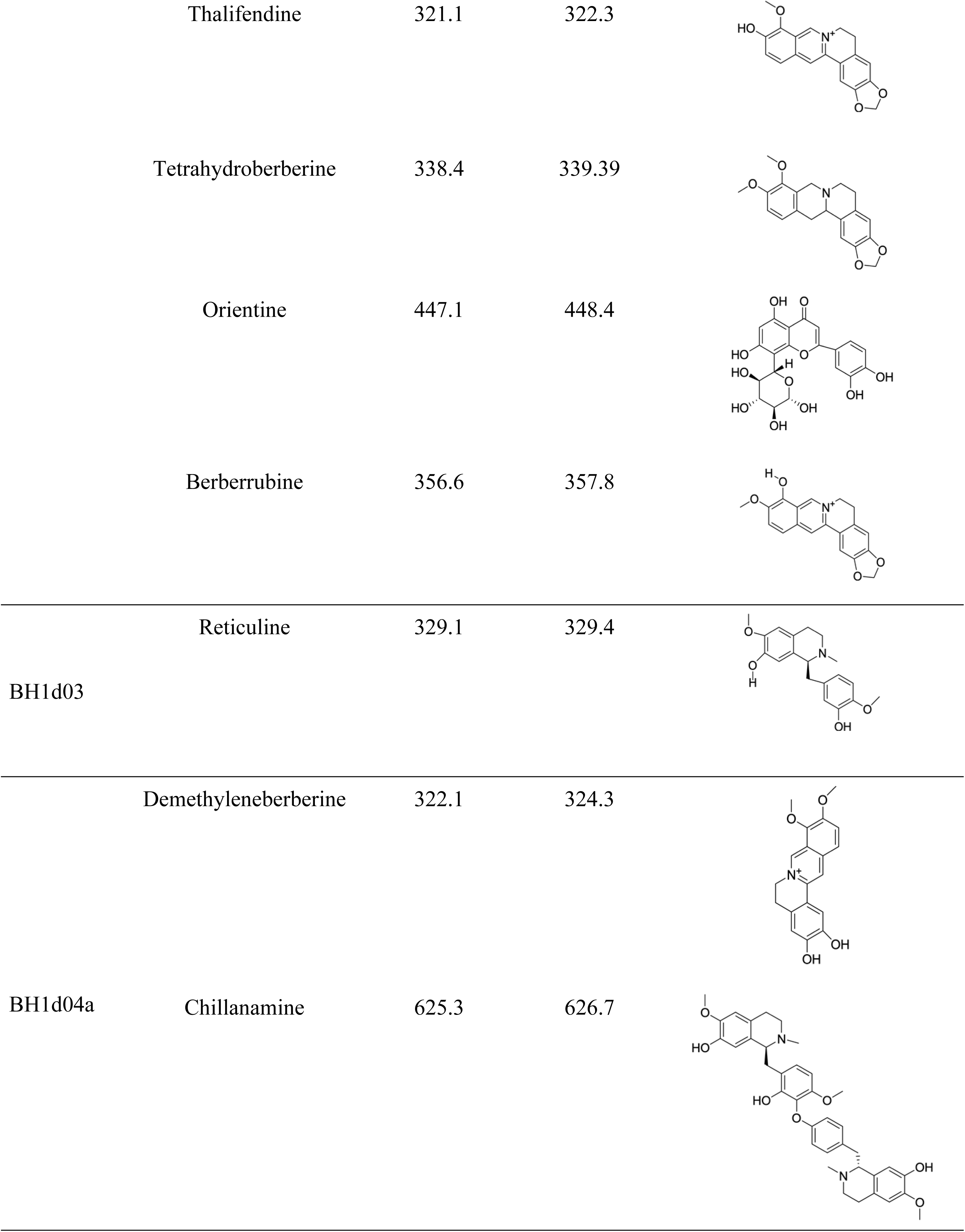
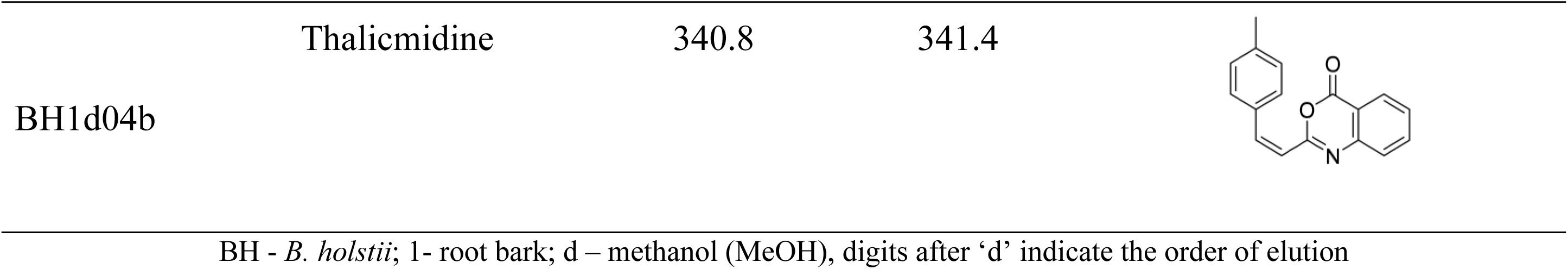
Compounds identified in the MeOH extract of the root bark of *B. holstii*.

To support our findings, the compounds identified in this study, in the root, stem, and leaf of other *Berberis* species have been reported in various published studies as shown in Table **6**.

**Table 6.** Compounds identified in *B. holstii* and reported in other *Berberis* species.

| Compound | Plant part | *Berberis* species | References |
| --- | --- | --- | --- |
| Thalcmidine | Aerial parts | *B. turcomanica* | (47) |
| Chillanamine | - | *B. buxifolia* | (48) |
| Isocorydine | Stem bark, leaves | *B. vulgaris* | (48) |
| Columbamine | Root bark, stem bark | *B. vulgaris*, *B. lamberti* | (46,49) |
| Berberrubine | Aerial parts | *B. sibirica* | (46,49) |
| Magnoflorine | Root bark | *B. lyceum*, *B. kansuensis* | (50,51) |
| Reticuline | Leaves, stem bark | *B. heterobotrys* | (46,52) |
| Pronuciferine | Aerial parts | *B. coletioides* | (46,52) |
| Orientine | - | *B. montana* | (53) |
| Tetrahydroberberine | Root bark, stem bark | *B. petiolaris* | (54) |
| Isoboldine | Aerial parts | *B. brandisiana* | (53,54) |
| Palmatine | Aerial part | *B. aristata* | (55,56) |
| Thalofoline | Root bark | *B. vulgaris* | (57) |
| Berberine | Aerial parts, root bark<br>stem bark | Many<br>*Berberis* species | (46) |
| Tetrahydropalmatine | Stem bark | *B. aristata* | (46,58) |
| N-methylcoclaurine | - | *B. montana* | (53) |

### 3.6 Checkerboard assays and *in silico* molecular docking of identified compounds

#### 3.6.1 Checkerboard assays of fractions and SPEC against WT *Msm*

After identifying the compounds within the bioactive fractions, and having demonstrated functionally the capacity of extracts to synergise the activity of conventional antimycobacterial drug (RIF), these fractions were subjected to checkerboard assays with SPEC against WT *Msm* (Table **7** and Figure **3**). SPEC was selected for this study due to its relatively narrow spectrum of activity, which is primarily attributed to efflux pump-mediated resistance mechanisms (23) in contrast to RIF, where resistance is predominantly caused by target site mutations (59–61). This characteristic makes SPEC a suitable candidate for investigation of efflux pump activity in *Mtb*.

**Figure 3:**
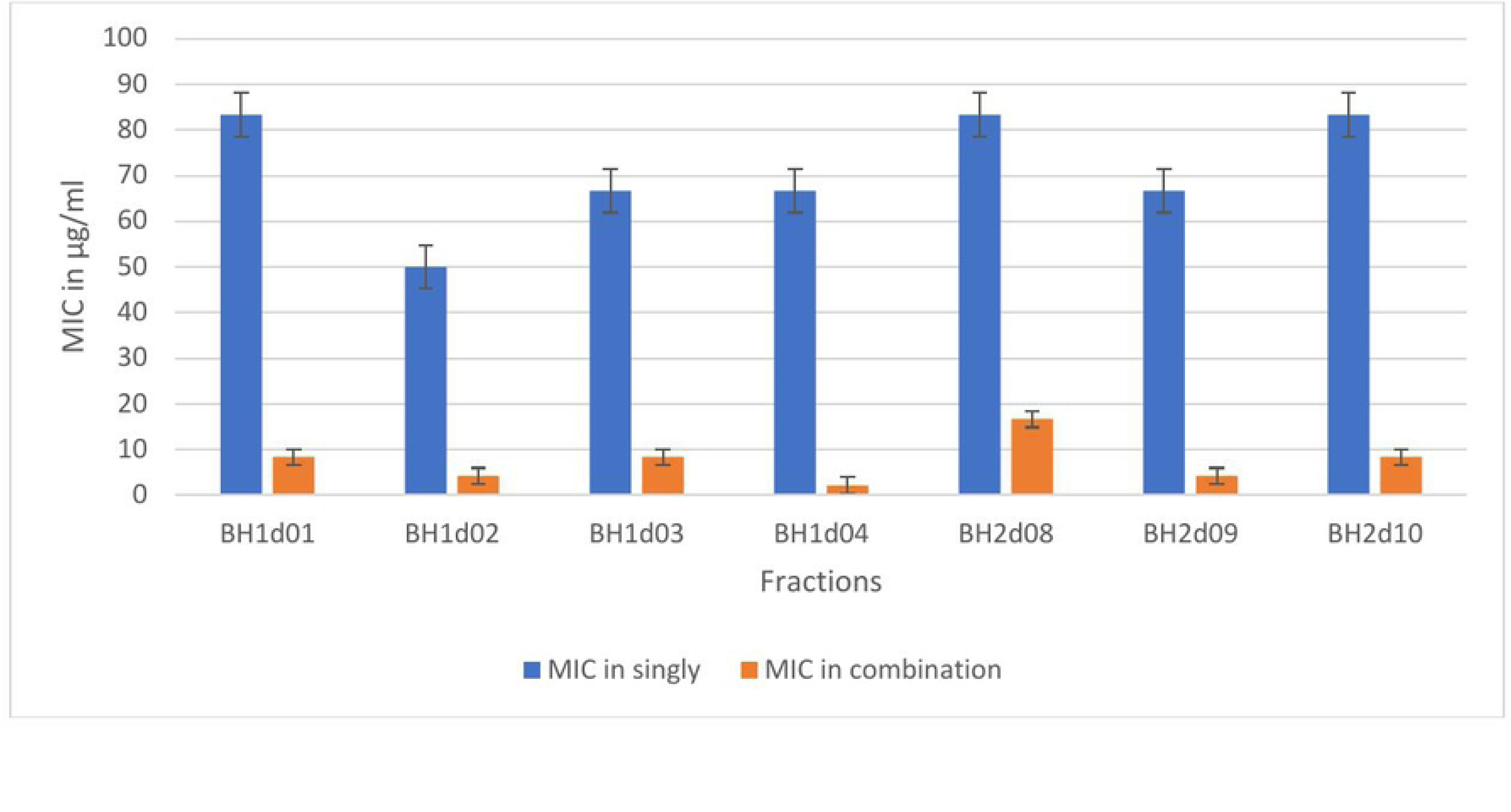
Checkerboard assay of SPEC with fractions against WT *Msm*; SPEC-spectinomycin; BH-*B. holstii*; 1-root bark; 2-stembark; d-methanol (MeOH), digits after ‘d’-order of elution

**Table 7.**
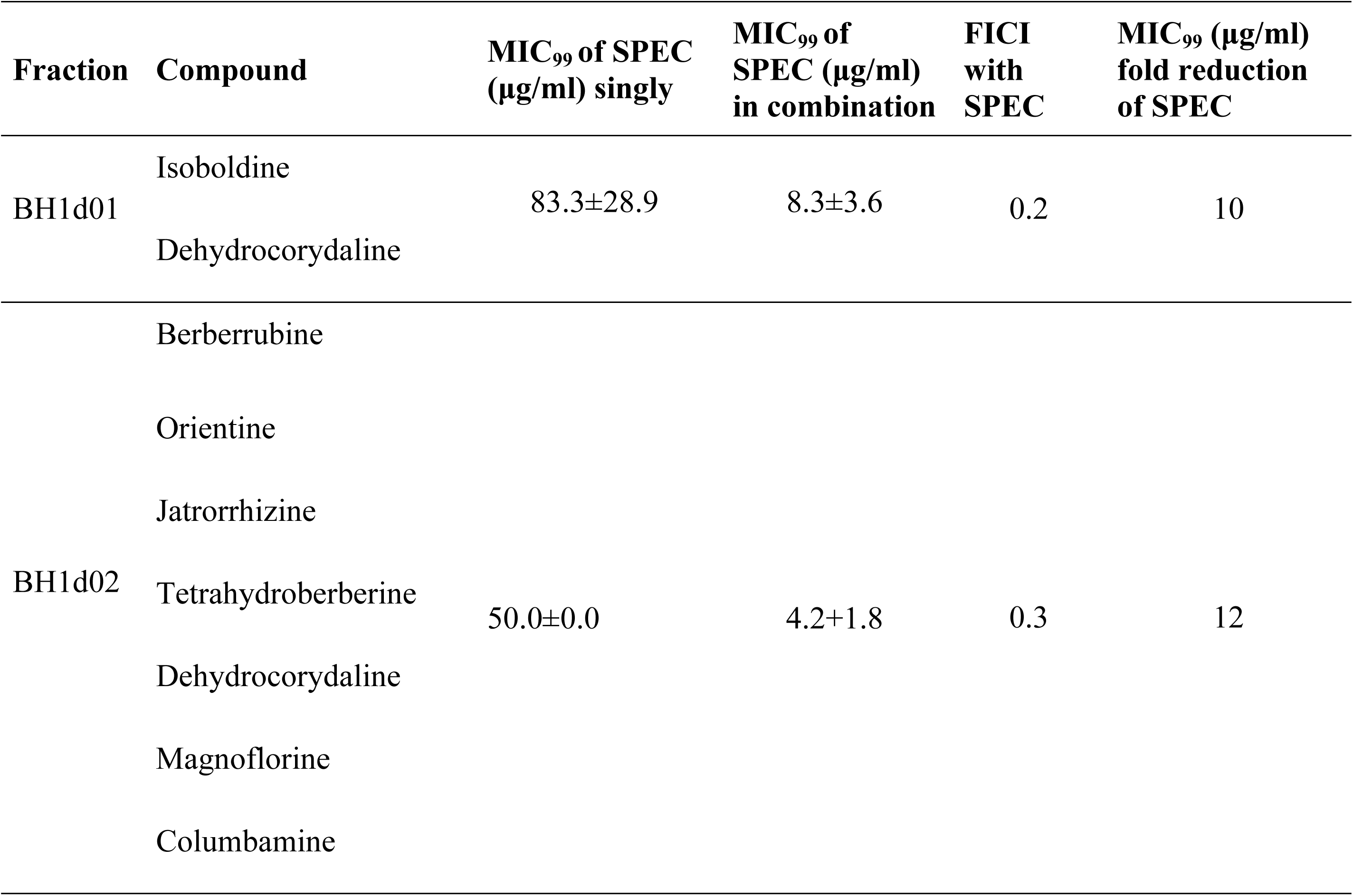
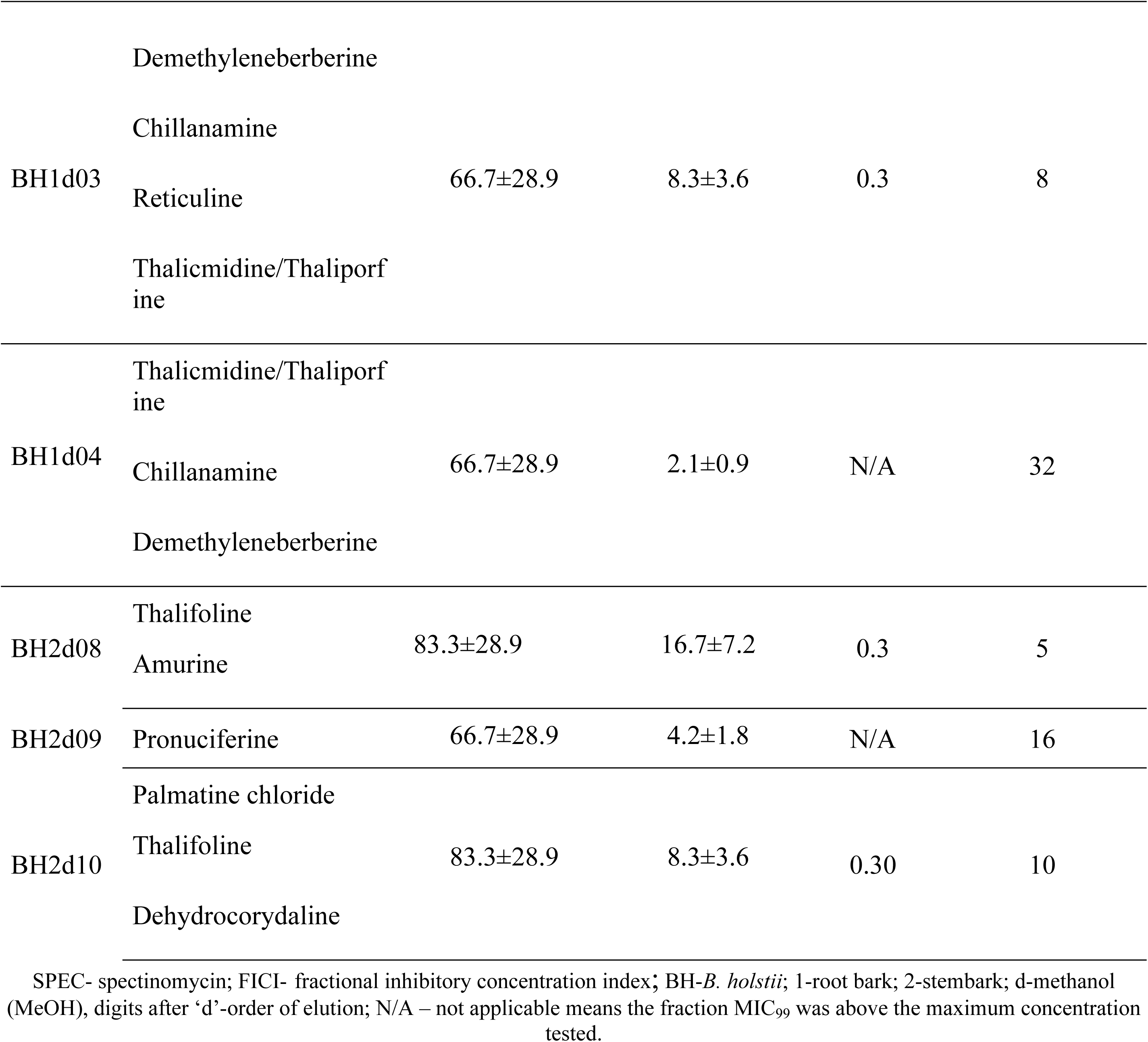
Checkerboard assay results for selected fractions of *B. holstii* and SPEC against WT *Msm*.

Five fractions exhibited FICI values below 0.4 with BH1d01 revealing the most potent interaction with FICI of 0.2 with SPEC against wild-type *Msm*. BH1d04 demonstrated the greatest enhancement of SPEC activity, resulting in a 32-fold reduction in its MIC₉₉ besides not exhibiting direct antimycobacterial activity against WT *Msm*.

#### 3.6.2 *In silico* molecular docking of identified compounds and Checkerboard assays of fractions and SPEC against WT *Mtb*

With promising results of Checkerboard assays against WT *Msm*, we sought to computationally infer compounds that could be associated with the demonstrated EP inhibition. For this, compounds identified by LC-MS in the bioactive fractions (Table **3**, Supplementary Table **S1** and Supplementary Table **S2**) were docked onto AlphaFold 3 generated three-dimensional models of Rv1258c, Rv2333c and MmpS5-MmpL5 *Mtb* EPs that have been implicated in the resistance to current anti-tubercular agents (12,62,63) to determine binding modes and affinities. The docking poses and scores are depicted in Figure **4** and Table **8** respectively Fractions containing compounds that exhibited docking scores of < -5 kcal/mol were further prioritised for screening in checkerboard assays with SPEC, an aminoglycoside antibiotic that is a substrate for mycobacterial Eps (64,65), against WT *Mtb* (Table **8** and Figure **5**).

**Figure 4.**
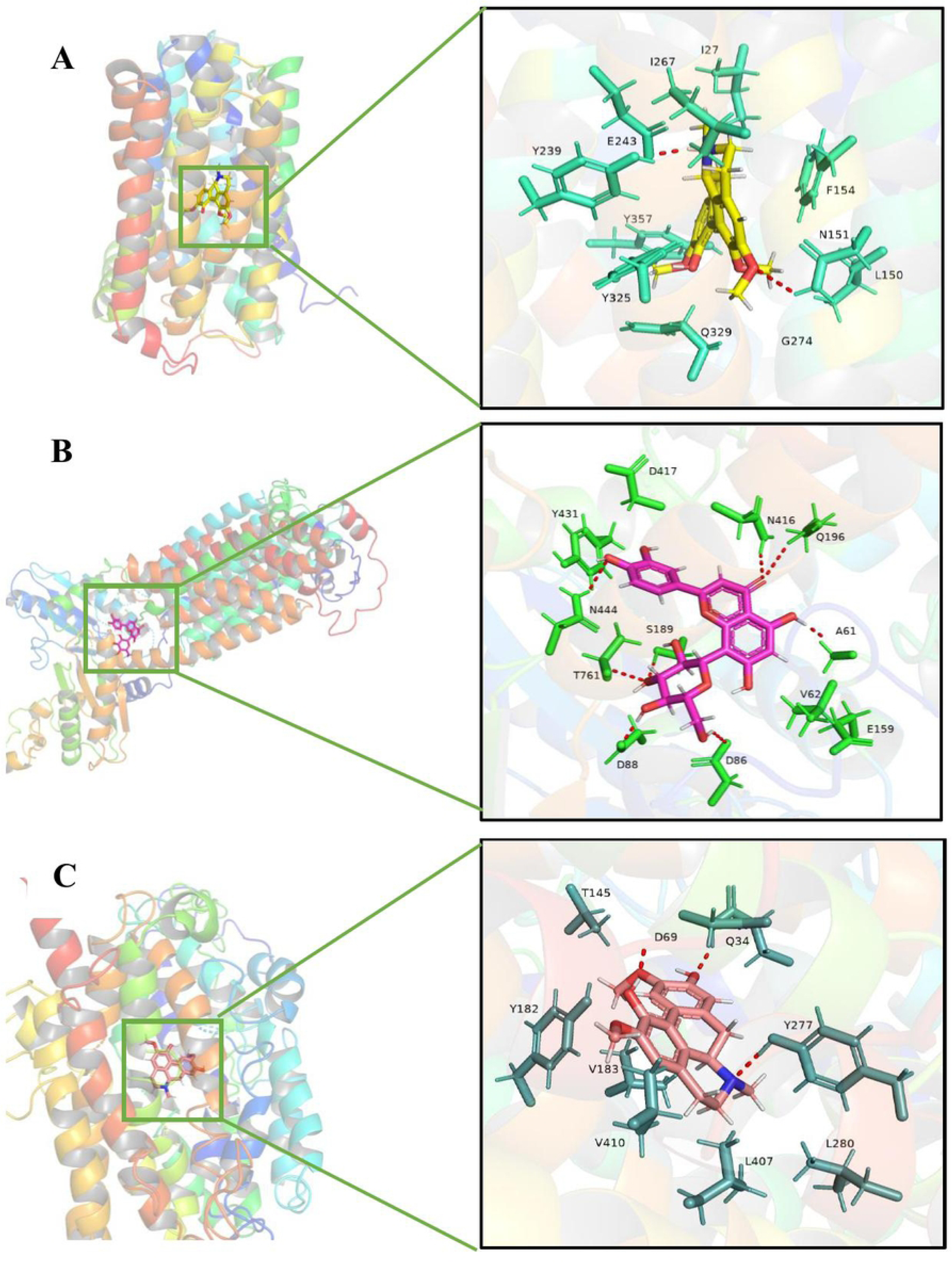
Molecular docking of select phytochemicals identified in *B. holstii* onto *Mtb* EPs. Highest scoring binding poses for isocorydine binding to Rv1258c, orientine binding to MmpS5-MmpL5 and isoboldine binding to Rv2333c are shown in panels A, B and C respectively.

**Figure 5:**
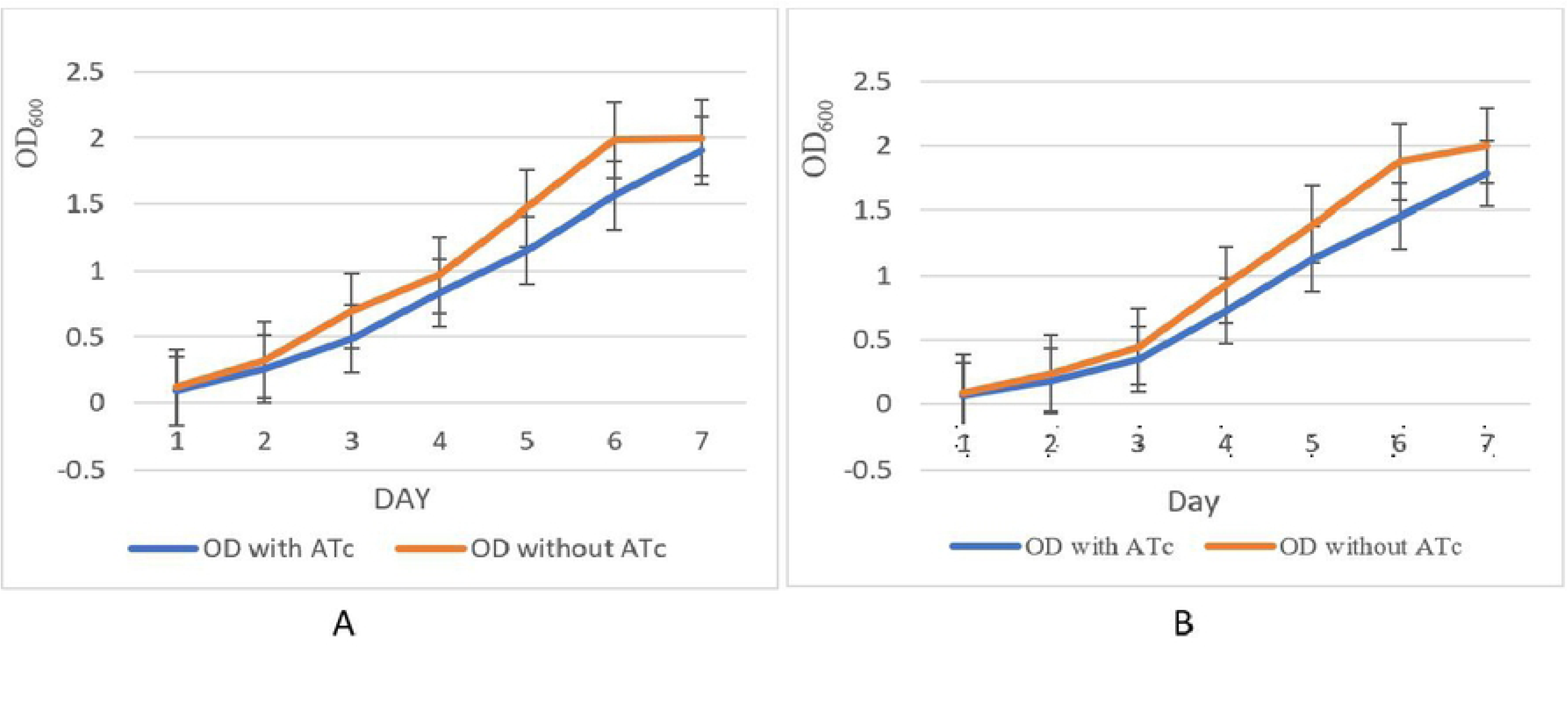
Checkerboard assays of SPEC with fractions against WT *Mtb;* SPEC-spectinomycin; BH-*B. holstii*; 1-root bark; 2-stembark; d-methanol (MeOH), digits after ‘d’-order of elution

**Table 8:**
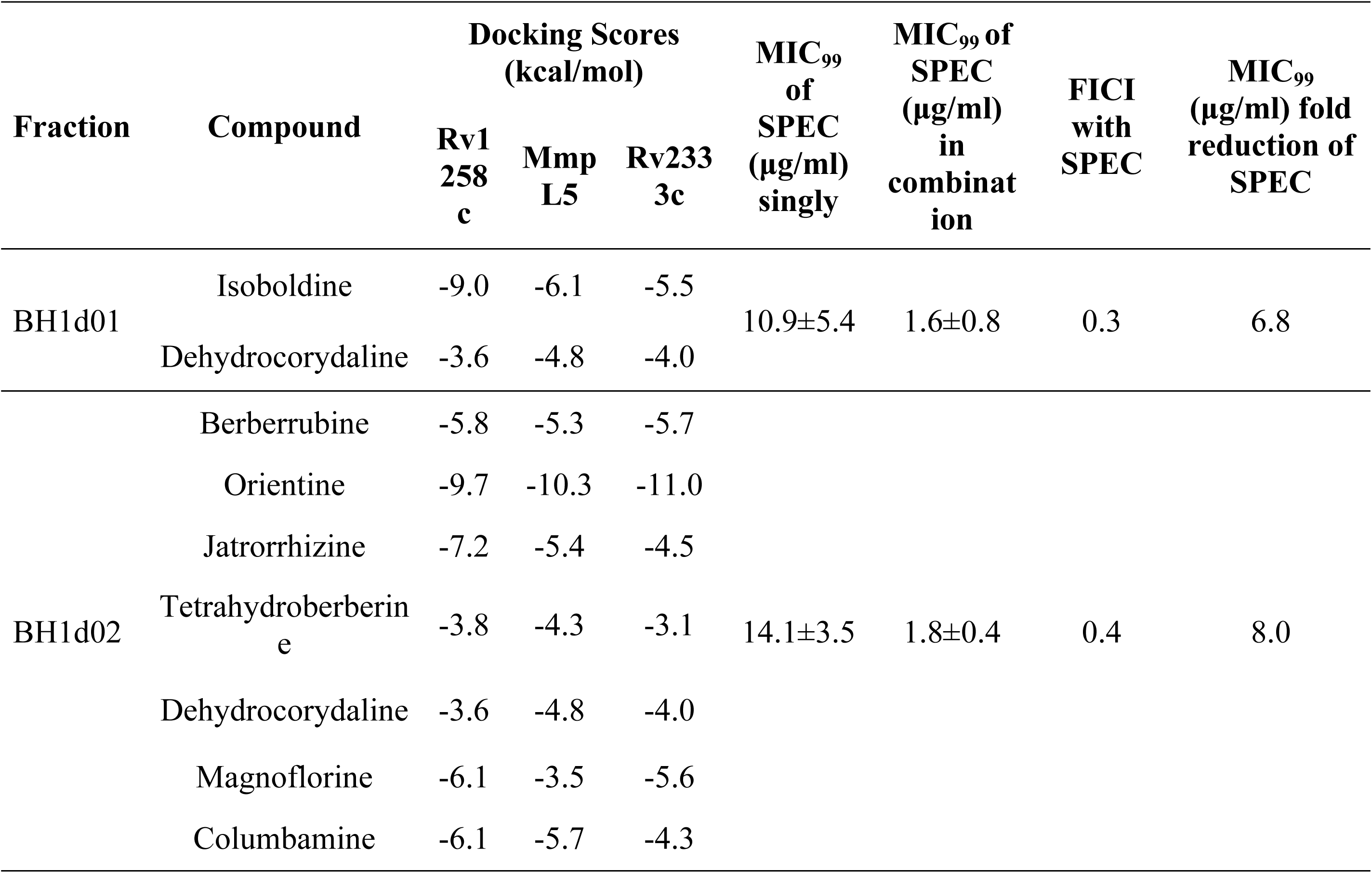
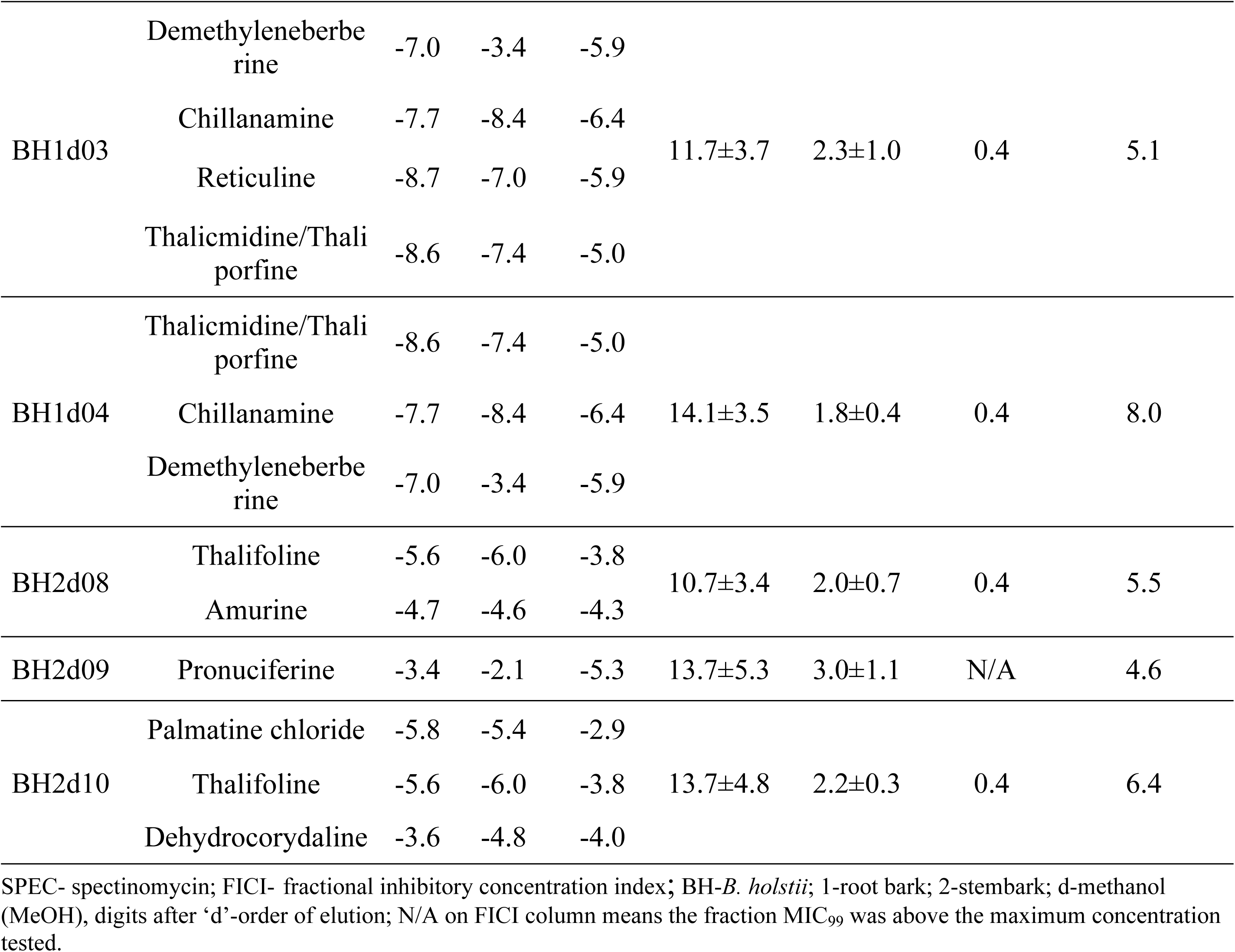
Checkerboard assay results for selected fractions of *B. holstii* and SPEC against WT *Mtb*.

Notably, only fractions deemed to contain compounds predicted to bind to *Mtb* Rv1258c, MmpS5-MmpL5 or Rv2333c were functionally assayed against WT *Mtb*. To confirm that the fractions were functioning as EPIs, they were screened against Rv1258c and Rv2333c CRISPRi knockdown *Mtb* strains and the growth fitness curves determined (Figure **6**) (Supplementary Table **S6,** Supplementary Table **S7)**. In both instances, the knockdown strains exhibited reduced growth rates in ATc-supplemented media compared to those cultured without ATc, consistent with the expected gene repression effect.

**Figure 6:**
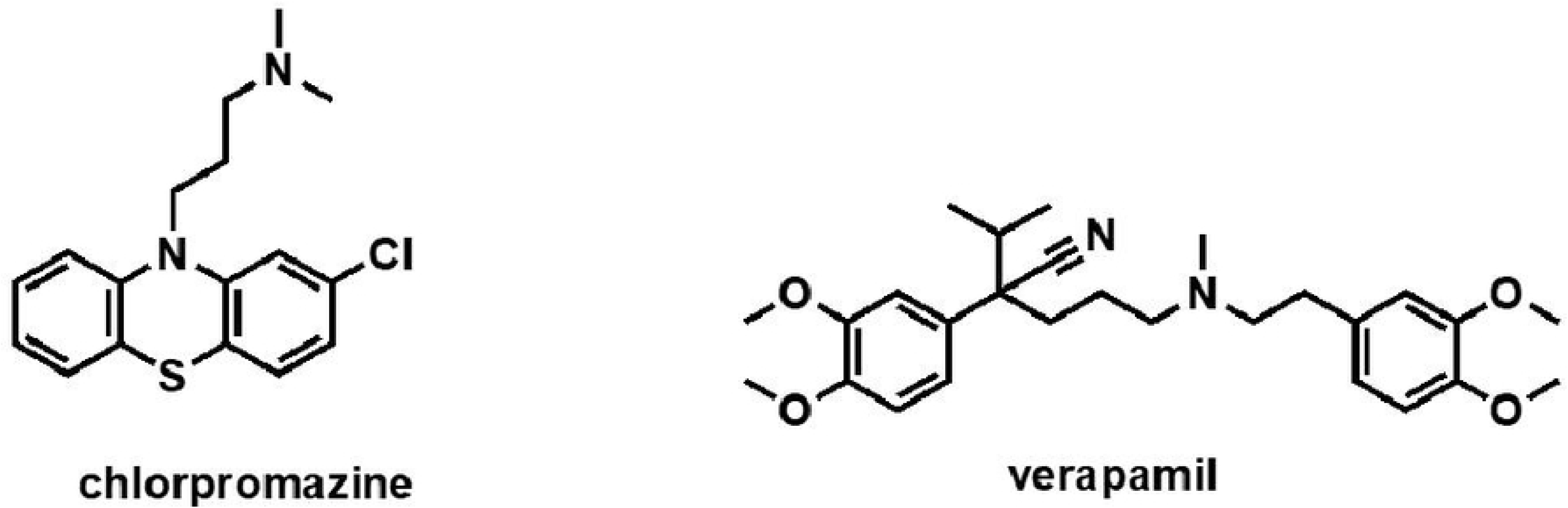
Growth curves of A, Rv1258c and B, Rv2333c *Mtb* CRISRPi knockdown strains in ATc and without ATc.

Then the MIC₉₉ of SPEC against H37Rv WT strain (15.63±5.41 μg/ml), Rv1258c knockdown (3.9±0.0 μg/ml) and Rv2333c knockdown (3.9±1.4 μg/ml) was determined (Table **9**). Notably both knockdown strains exhibited lower MIC₉₉ values compared to WT *Mtb* strain indicating increased susceptibility to SPEC upon efflux pump gene repression. Additionally, the MIC₉₉ of SPEC (1.6±0.8, 1.8±0.4, 2.3±1.0, 1.8±0.4, 2.0±0.7, 3.0±1.1, 2.2±0.3 μg/ml) in combination with the fractions (BH1d01, BH1d02, BH1d03, BH1d04, BH2d08, BH2d09, BH2d10 respectively) against WT *Mtb* was reduced (Table **8**) to values (3.9±0.0, 3.9±1.4 μg/ml) that are comparable to the MIC₉₉ values seen in Rv1258c and Rv2333c knockdown strains respectively (Table **9**).

**Table 9.** MIC assays of extract fractions with SPEC against WT *Mtb*, Rv1258c and Rv2333c knock down strains.

| Strain | H37Rv WT | Rv1258c | Rv2333c |
| --- | --- | --- | --- |
| MIC<sub>99</sub> in µg/ml | 15.6±5.4 | 3.9±0.0 | 3.9±1.4 |

## 4.0 Discussion

Overall, this study sought to ascertain whether the secondary metabolites of *B. holstii* have the ability to hamper the *Mtb* efflux process by blocking EP components that can act as potentiators of anti-TB drugs to which resistance has been reported. However, on assaying for antimycobacterial activity using WT *Msm*, the extracts’ activity was low compared to that of RIF (Table **2**). This may be attributed to dwindled quantity of bioactive compound(s) in the extracts or masking of the activities of the compound(s) by other metabolites (66). Although the total MeOH extract from the leaves revealed relatively high activity against WT *Mtb* (MIC_99_ of 60.43 μg/ml), the extract showed low activity against WT *Msm* (MIC_99_ of 4500 μg/ml) (Table **2**). Compared to other total extracts, the aqueous extract of the stem bark was highly active against WT *Mtb* (MIC_99_ of 1.738 μg/ml) (Table **2**) even though this activity was lower than that of the RIF positive control. Moreover, the MeOH and aqueous extracts from the roots displayed relatively minimized activity against WT *Msm* (MIC_99_ values of 906.25 μg/ml and 1875.00 μg/ml respectively) (Table **2**) compared to the RIF positive control. The modest activity of the plant extracts against WT *Msm* and WT *Mtb* plausibly reflects the low total quantity of bioactive compounds in each sample (67). This observation may explain the fact that though *B. holstii* has been reported to possess EP inhibitory activity (67), no studies have reported it to possess mycobacteria inhibitory activity. The complement of *B. holstii* extracts analyzed showed tolerable cytotoxicity against Vero cells that is within the National Cancer Institute weakly cytotoxic category (CC_50_<20 μg/mL highly cytotoxic, CC_50_ 21 to 200 μg/mL moderately cytotoxic, CC_50_ 201 to 500μg/mL weakly cytotoxic, and CC_50_>500 μg/mL noncytotoxic) (42–45).

Checkerboard assay results indicated that the aqueous and MeOH total extracts were synergistic with RIF which corroborates the study that reported *B. holstii* to contain compounds with probable *Mtb* EP inhibitory activity (67). On the other hand, DCM extracts of the stem bark exhibited an additive effect. Total extracts delineated to contain potential *Mtb* EPs blocking activity were thereafter fractionated using column chromatography and the constituent compounds uncovered using LC-MS. Specifically, compound identification was based on molar masses and fragmentation patterns of individual compounds as compared to those reported in literature and the NLM, Lotus and GNPS databases (53,54,57).

Molecular docking of compounds identified using LC-MS onto *Mtb* Rv1258c, Rv2333c and MmpS5-MmpL5 EP proteins that have been implicated in resistance to current anti-tubercular agents (12,62,63) revealed moderate to strong binding affinities. Subsequently, extract fractions containing compounds with robust docking scores were screened against WT *Msm* in combination with SPEC, an aminoglycoside indicated for the treatment of uncomplicated gonorrhoeae (68) that is extruded by mycobacteria EPs (64,65). The very low antimycobacterial activity of SPEC was notably enhanced (>100 fold) by EPI such as chlorpromazine and verapamil (Figure **7**) (28,69). Analogues of SPEC known as spectinamides have been shown to inhibit EPs resulting in high *in vitro* activity against *Mtb* (23). However, these compounds do not exhibit *in vivo* antimycobacterial activity.

**Figure 7:**
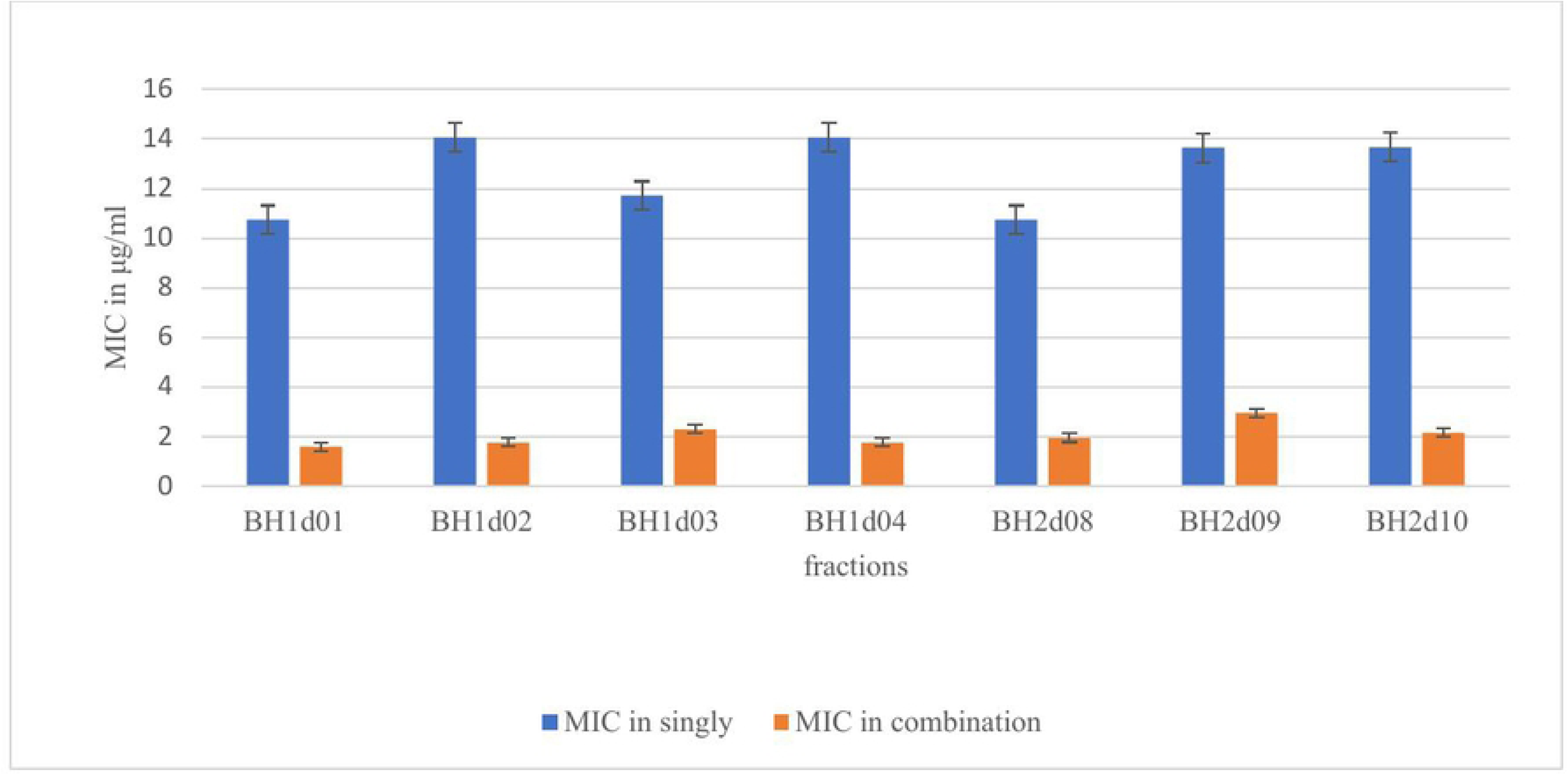
Examples of known EPIs

Antimycobacterial bioassays and checkerboard assays conducted on 10 individual fractions showed that six exhibited low activity against WT *Msm* (MIC₉₉ ranging from 150 to 860 μg/ml) compared to RIF (MIC₉₉ 12.75 μg/ml), while the other six demonstrated moderate activity against WT *Mtb* (MIC₉₉ ranging from 54.7 to 287.5 μg/ml) relative to SPEC (MIC₉₉ 18.8 μg/ml) (supplementary Table **S3**). The BH1d01 fraction, which was the first to elute during chromatographic separation of the MeOH root bark extract, exhibited one of the strongest *in vitro* synergistic effects—showing a FICI of 0.2 (Table **7**) against WT *Msm* and 0.3 (Table **8**) in combination with SPEC against WT *Mtb*.

Notably, LC-MS analysis revealed the fraction to contain isoboldine (Table **5**), a compound predicted to bind to Rv1258c, MmpS5-MmpL5 and Rv2333c with strong affinity (-9.0, -6.1, - 5.5kcal/mol, respectively) Table **8**. Isoboldine, previously identified in the stem bark extracts of *Annona cherimolia* (71) has been shown to possess antibacterial properties(72). The other major constituent uncovered in BH1d01, dehydrocorydaline, had predicted weak binding to the *Mtb* efflux proteins and is therefore unlikely to contribute to any observed synergistic activity of this fraction. Another fraction that exhibited high *in vitro* synergistic effect was BH2d10 (FICI=0.30) with WT *Msm* but relatively lower with WT *Mtb* (FICI=0.4) Table **8** and Supplementary Table **S5**. This fraction was ascertained to contain palmatine chloride, dehydrocorydaline, and thalifoline. Palmatine chloride and thalifoline Table **8** were predicted to bind to Rv1258c and MmpS5-MmpL5 with moderate affinity and may therefore be contributors to the observed EI. Palmatine has been shown to block EPs in several antibiotic resistant bacteria including *Aeromonas hydrophila* and *Pseudomonas aeruginosa* (73–75). The BH1d03 fraction ascertained to contain thalicmidine, reticuline, chillanamine and demethyleneberberine also exhibited *in vitro* synergistic activity against WT *Msm* (FICI=0.3) and WT *Mtb* (FICI=0.4) Table **8** and Supplementary Table **S5**. Molecular docking predicted chillanamine and reticuline to strongly bind to the three EPs evaluated. Specifically, thalicminide was predicted to strongly bind (-8.6, -7.4kcal/mol respectively) to Rv1258c and MmpS5-MmpL5, while demethyleneberberine (-7.0, -5.9kcal/mol respectively) bound to Rv1258c and Rv2333c (Table **8**). These *in vitro* and *in silico* data, point to demethyleneberberine, a natural analogue of berberine, being a presumptive EPI. Berberine is a natural compound that has shown plausible EP inhibition activity in MDR bacterial isolates (73–76). Nevertheless, reticuline has been reported to be inactive against several bacteria species including *P. aeruginosa*, *Escherichia coli*, *Bacillus cereus*, *B. subtilis* and *Staphylococcus aureus* (77,78).

An additional fraction, BH2d08, containing thalifoline and amurine, also demonstrated moderate synergistic activity, with a FICI of 0.3 against WT *Msm* (Table **7** Supplementary Table **S4**) and 0.4 against WT *Mtb* (Table **8** Supplementary Table **S5**). The two compounds contained in the fraction were predicted to bind to Rv1258c and MmpS5-MmpL5, and may therefore be responsible for the synergistic effect of the fraction. Thalifoline has been reported to exhibit antifungal activity against *Fusarium oxysporum* (79). The final fraction that exhibited *in vitro* synergistic activity with both WT *Msm* (FICI = 0.3) (Table **7** Supplementary Table **S4**) and WT *Mtb* (FICI = 0.4) (Table **8** Supplementary Table **S5**) was BH1d02, which contained berberrubine, orinetine, jatrorrhizine, tetrahydroberberine, dehydrocorydaline, magnoflorine, and columbamine. Based on *in silico* docking analyses, this synergistic activity may be attributed to berberrubine, orientine, jatorrhizine, magnoflorine and columbamine which all manifested strong binding affinities onto the EPs evaluated. Berberrubine, a primary active metabolite of berberine, has been reported as a likely inhibitor of mycobacterial EPs (80). Akin to berberine, columbamine and tetrahydroberberine are protoberberine alkaloids with reported antibacterial activity against *E. coli*, *S. aureus*, *B. subtilis*, *Salmonella enterica* serovar Enteritidis, *Helicobacter pylori* and *Brucella abortus* (81–83). Consistent with our results, jatrorrhizine has previously been reported to be an inhibitor of bacterial EPs (84). Besides, the *Ocimum sanctum* flavonoids orientine and vicenin have been demonstrated to synergistically inhibit the growth of *E. coli, S. aureus*, *K. pneumoniae*, *Staphylococcus cohnii* and *Proteus* (85). Although our *in silico* studies did not reveal dehydocorydaline to robustly bind to the EPs, Kim and co-workers have reported that it has strong antibacterial activity against *E. coli*, *S. enterica* serovar Enteritidis, *S. aureus* and *Listeria monocytogenes*, with the activity analogous to that of berberine (83).

Interestingly, the BH2d09 fraction did not exhibit direct antimycobacterial activity against either WT *Msm* or WT *Mtb*, yet it significantly enhanced the efficacy of SPEC reducing its MIC₉₉ by 16-fold (Table **7**, Supplementary Table **S4**) against WT *Msm* and by 4.6-fold (Table **8**, Figure **5**, Supplementary Table **S5**) against WT *Mtb*. The only compound identified in this fraction was pronuciferine, a naturally-occurring quinolone. A similar effect was observed for the BH1d04 fraction that remarkably (32-fold decrease) (Table **7**, Figure **4**, Supplementary Table **S4**) diminished SPEC MIC_99_ devoid of antimycobacterial activity against WT *Msm* however this particular fraction exhibited synergistic activity against WT *Mtb* with FICI of 0.4 (Table **8**, Supplementary Table **S5**) with SPEC. This fraction contains thalicmidine, chillanamine and demethyleneberberine that are predicted to strongly bind to Rv1258c, MmpS5-MmpL5 and Rv2333c. Importantly, in the context of EI, compounds do not necessarily have to exhibit a killing effect on an organism, however, their ability to block the extrusion of the active partner enhances the overall biological activity as observed here and in similar reported studies (28,86). To elucidate the potential mechanism of action of the fractions, CRISPRi knockdown strains Rv1258c and Rv2333c were employed. These strains exhibited a two- to three-fold reduction in the MIC₉₉ of SPEC relative to the H37Rv wild-type strain (Table **9**). Furthermore, the MIC₉₉ of SPEC in combination with the fractions against the WT *Mtb* strain (Table **8**) was comparable to the MIC₉₉ of SPEC against the knockdown strains (Table **9**), suggesting that the compounds in the fractions may be functioning as EPIs.

In sum, our computational and experimental results have delineated *B. holstii* compounds with probable mycobacterial EP inhibitory activity namely isoboldine, berberrubine, orientine, jatrorrhizine, magnoflorine, columbamine, demethyleneberberine, chillanamine, reticuline, thalicmidine, thalifoline and palmatine chloride. Importantly, this study has demonstrated the utility of integrating computational methods with traditional natural product screening approaches for the identification of putative mycobacterial EPIs. Notably, while the antibacterial activity of berberine, palmatine chloride, columbamine and jatrorrhizine has previously been reported, to the best of our knowledge, ours is the first study to investigate these compounds as putative inhibitors of *Mtb* EPs with the potential to augment the efficacy of current TB drugs by mitigating resistance and consequently shortening the lengthy therapy duration (28). The compounds identified in this study warrant further investigation as potential lead entities in the TB drug development pipeline.

## 5.0 Conclusion

In this study, we have uncovered *B. holstii* compounds with potential *Mtb* EP inhibitory activity that may be further evaluated as leads for buttressing the activity of current anti-TB agents to which resistance due to extrusion has been reported. Our findings suggest that *B. holstii* derived compounds might block SPEC efflux and thereby potentiate the antimycobacterial activity of anti-TB compounds. This study is a proof of concept that plant derived compounds may be paired with current anti-TB drugs and those in development to circumvent the development and propagation of resistance underpinned by drug extrusion. Further investigation of the identified compounds is warranted.

## Acknowledgments

We express our gratitude to The World Academy of Sciences for their support through Grant No. RG/CHE/AF/AC_I – FR3240297746, The Grand Challenges-Africa Program Round 10-2nd Drug Discovery for support via Grant No. GCA/DD2/Round10/022/002, the Medical Research Foundation for funding via Grant No. MRF-131-0005-RG-RACE-C0853 and BBSRC (BB/M025624/1). We extend our gratitude to the late Dr. Bernald Wanjohi from the University of Eldoret, a botanist who contributed to plant collection and identification, and to Mr. Nicolus Adipo from CTMDR, KEMRI, for his assistance in conducting cytotoxicity assays. Additionally, we thank to Dr. Peter Wilson from the School of Biochemistry at the University of Bristol for his technical contributions during LC-MS analysis.

